# Niche and neutral processes leave distinct structural imprints on indirect interactions in mutualistic networks

**DOI:** 10.1101/2020.04.30.070391

**Authors:** Benno I. Simmons, Andrew P. Beckerman, Katrine Hansen, Pietro K. Maruyama, Constantinos Televantos, Jeferson Vizentin-Bugoni, Bo Dalsgaard

## Abstract

Indirect interactions are central to ecological and evolutionary dynamics in pollination communities, yet we have little understanding about the processes determining patterns of indirect interactions, such as those between pollinators through shared flowering plants. Instead, research has concentrated on the processes responsible for direct interactions and whole-network structures. This is partly due to a lack of appropriate tools for characterising indirect interaction structures, because traditional network metrics discard much of this information. The recent development of tools for counting motifs (subnetworks depicting interactions between a small number of species) in bipartite networks enable detailed analysis of indirect interaction patterns. Here we generate plant-hummingbird pollination networks based on three major assembly processes – neutral effects (species interacting in proportion to abundance), morphological matching and phenological overlap – and evaluate the motifs associated with each one. We find that different processes produce networks with significantly different patterns of indirect interactions. Neutral effects tend to produce densely-connected motifs, with short indirect interaction chains, and motifs where many specialists interact indirectly through a single generalist. Conversely, niche-based processes (morphology and phenology) produced motifs with a core of interacting generalists, supported by peripheral specialists. These results have important implications for understanding the processes determining indirect interaction structures.

## Introduction

Species in a community are often influenced by other species they do not interact with directly (Strauss, 1991; Wootton, 1994, 2002). Such indirect interactions are a fundamental component of communities, governing ecological and evolutionary processes as much as, or more than, direct effects (Bailey & Whitham, 2007; Guimarães, Pires, Jordano, Bascompte, & Thompson, 2017; Martínez, García, & Herrera, 2014; Strauss, 1991; Vandermeer, Hazlett, & Rathcke, 1985). For example, in plant-pollinator communities, indirect interactions between plants can be mediated by shared pollinator species. These can be facilitative, where one plant attracts pollinators that also visit co-occurring plant species, or competitive, where one plant attracts pollinators away from another plant, through being more abundant or more attractive to the pollinator than the competing plant (Carvalheiro et al., 2014; Mitchell, Flanagan, Brown, Waser, & Karron, 2009; Morales & Traveset, 2009). These indirect interactions can have important implications for community persistence and stability. For example, in communities dominated by apparent competition, the sharing of interaction partners is restricted, and thus perturbations are limited in how much they can propagate through the community (Thébault & Fontaine, 2010). Conversely, communities dominated by apparent facilitation favour connected, nested structures with enhanced species coexistence (Bastolla et al., 2009; Thébault & Fontaine, 2010).

Despite the importance of indirect interactions for ecological and evolutionary dynamics, we have little understanding of the processes that lead to their formation and maintenance in mutualistic networks. Instead, research has focused on determining processes that give rise to whole-network patterns or individual direct interactions, leaving the determinants of local-scale patterns of indirect interactions largely unexplored (Maruyama, Vizentin-Bugoni, Oliveira, Oliveira, & Dalsgaard, 2014; Olito & Fox, 2015; Simmons, Vizentin-Bugoni, et al., 2019; Vázquez, Chacoff, & Cagnolo, 2009).

Knowledge of the processes responsible for indirect interactions is not only important in terms of understanding the assembly and maintenance of community structure, but could also have implications for conservation. Three distinct processes have been used to explain mutualistic network structure: morphological matching (similarity in size and shape of a flower’s corolla and a pollinators feeding apparatus [Sonne et al., 2020b]); phenological overlap (co-occurrence in time of a flower and pollinator), and neutral effects (assembly based on species interacting randomly in proportion to their abundance). If, for example, indirect interaction structures are a result of neutral effects, then conservation might focus on preserving species’ abundance distributions. If network structure is primarily determined by morphological matches between species, then conservation might focus on ensuring the presence and persistence of species with complementary sets of morphological traits. If phenological overlap between species is the main process that governs network structure, then conservation might need to ensure the community comprises sets of species with synchronous timings, so that interactions remain established under climate change. Moreover, if it can be established that different processes form different indirect interaction structures – that is, if different processes leave distinct structural imprints – it may be possible to infer the processes operating in a community from network topology alone, without having to expend valuable time and money collecting the extra data required to measure processes explicitly.

Here we aim to understand the determinants of different indirect interaction structures by comparing the indirect interaction structures produced by three distinct assembly processes: morphological matching and phenological overlap (collectively known as niche-based processes), and abundance (neutral effects). We use 24 empirical datasets on species abundance, morphology and phenology from plant-hummingbird pollination communities across the Americas to understand network structures that result from different processes. We find that different processes leave distinct imprints on the structure of indirect interactions in mutualistic networks, and conclude that this could have important implications for conservation in the future.

## Materials and Methods

We created simulated networks under three processes (morphological matching, phenological overlap and neutral effects) from a dataset of 24 plant-hummingbird pollination networks (Sonne et al., 2020b, 2020a), which contained corresponding information on plant and hummingbird abundance, morphology (hummingbird bill length and floral corolla depth) and phenology. Communities sampled span from Mexico to Brazil. Full details of the data are given in Sonne et al. (2020b).

For each of these sets of abundance, morphology and phenology data, we generated matrices giving the probabilities of species interactions under three different processes, following Vázquez *et al.* (2009): neutral effects, morphological matching and phenological overlap. Neutrality was simulated using an abundance matrix, **A**. Elements of **A** were the product of each species’ relative abundance. Thus, element *a*_*ij*_ represents the interaction probability between plant species *i* and hummingbird species *j* and is equal to the product of the relative abundances of *i* and *j*. This matrix therefore represents neutrality: the likelihood of species interacting randomly in proportion to their abundance.

We create two morphological match matrices, corresponding to two different methods in the literature. In the first matrix, **M**_**F**_, hummingbird bill lengths were first multiplied by 4/3 to account for the extension of the tongue beyond the length of the bill (J. Vizentin-Bugoni, Maruyama, & Sazima, 2014). Matrix elements were then set to 1 if the bill length (plus the extension of the tongue) equalled or exceeded the floral corolla depth, and 0 otherwise (J. Vizentin-Bugoni et al., 2014). This follows the ‘forbidden link’ concept where species are only able to interact if there is a morphological match (i.e. if the hummingbird can reach the nectar in the floral corolla). Matrix elements were then divided by the sum of the matrix to convert the elements to probabilities (J. Vizentin-Bugoni et al., 2014). In the second matrix, **M**_**D**_, probabilities were inversely proportional to the difference between floral corolla depth and hummingbird bill length (Weinstein & Graham, 2017). This approach relaxes the assumption that a hummingbird is equally likely to interact with all flowers that have a floral corolla equal to or shorter than its bill, and makes morphological match a continuous, rather than binary, quantity. If the difference between floral corolla depth and hummingbird bill length was 0, the difference was set to the minimum non-zero difference between corolla depth and bill length in the web to prevent errors when dividing by zero values.

Elements of the phenological overlap matrix, **P**, were calculated using matrix multiplication (Vázquez et al., 2009). Plant and hummingbird phenology data, **O**_**P**_ and **O**_**H**_ respectively, had species as rows and dates as columns, with cells set to 1 for presence and 0 for absence of hummingbirds/flowers. Phenological overlap was then quantified as **P** = **O**_**P**_ **O**_**H**_′, where ′ indicates the matrix was transposed (Vázquez et al., 2009). Thus, element *p*_*ij*_ of **P** represents the number of time slices in which plant species *i* and hummingbird species *j* co-occur.

For each assembly process, and for each dataset, we generated 1000 binary interaction matrices from the probability matrix using the ‘mgen’ function in the ‘bipartite’ R package (Dormann, Frund, Bluthgen, & Gruber, 2009). In total there were 96,000 binary matrices (1000 generated matrices × four assembly processes × 24 sets of abundance, morphology and phenology data). Generated matrices had the same connectance as their corresponding empirical matrices.

### Characterising indirect interactions using motifs

We next characterised the different patterns of indirect interactions for each network and assembly process. Mutualistic networks are generally characterised using metrics that capture a particular facet of whole-network structure in a single number, such as levels of connectance, nestedness and modularity (Dalsgaard et al., 2013; Olesen & Jordano, 2002). While these metrics are undoubtedly useful, they are not always appropriate for considering indirect interactions in detail because compressing a network into a single number necessarily discards a substantial amount of topological information about indirect interactions (Simmons, Cirtwill, et al., 2019).

Here we instead characterise network structure using motifs, which have recently been proposed as a way to capture indirect interactions in bipartite networks in much greater detail than traditional metrics like nestedness and modularity (Simmons, Cirtwill, et al., 2019; Simmons, Sweering, et al., 2019). As motifs are a relatively new technique in the study of mutualistic networks, below we provide a brief introduction to the approach and outline a motif typology to aid their interpretation.

Just as LEGO sets are complex structures made from many small, distinct parts (Jordano, 2016a), networks can be thought of as being composed of many small subnetworks, or ‘building blocks’, known as motifs. Motifs take the form of small groups of species interacting with each other in particular ways. While network-level metrics like connectance, nestedness and modularity characterise network structure at the ‘macro-scale’, and species-level metrics like degree, *d*’, and centrality measures characterise the role of individual nodes at the ‘micro-scale’, motifs sit between these two extremes and capture ‘meso-scale’ network structure: local patterns of indirect interactions (Figure 1a).

**Figure 1:**
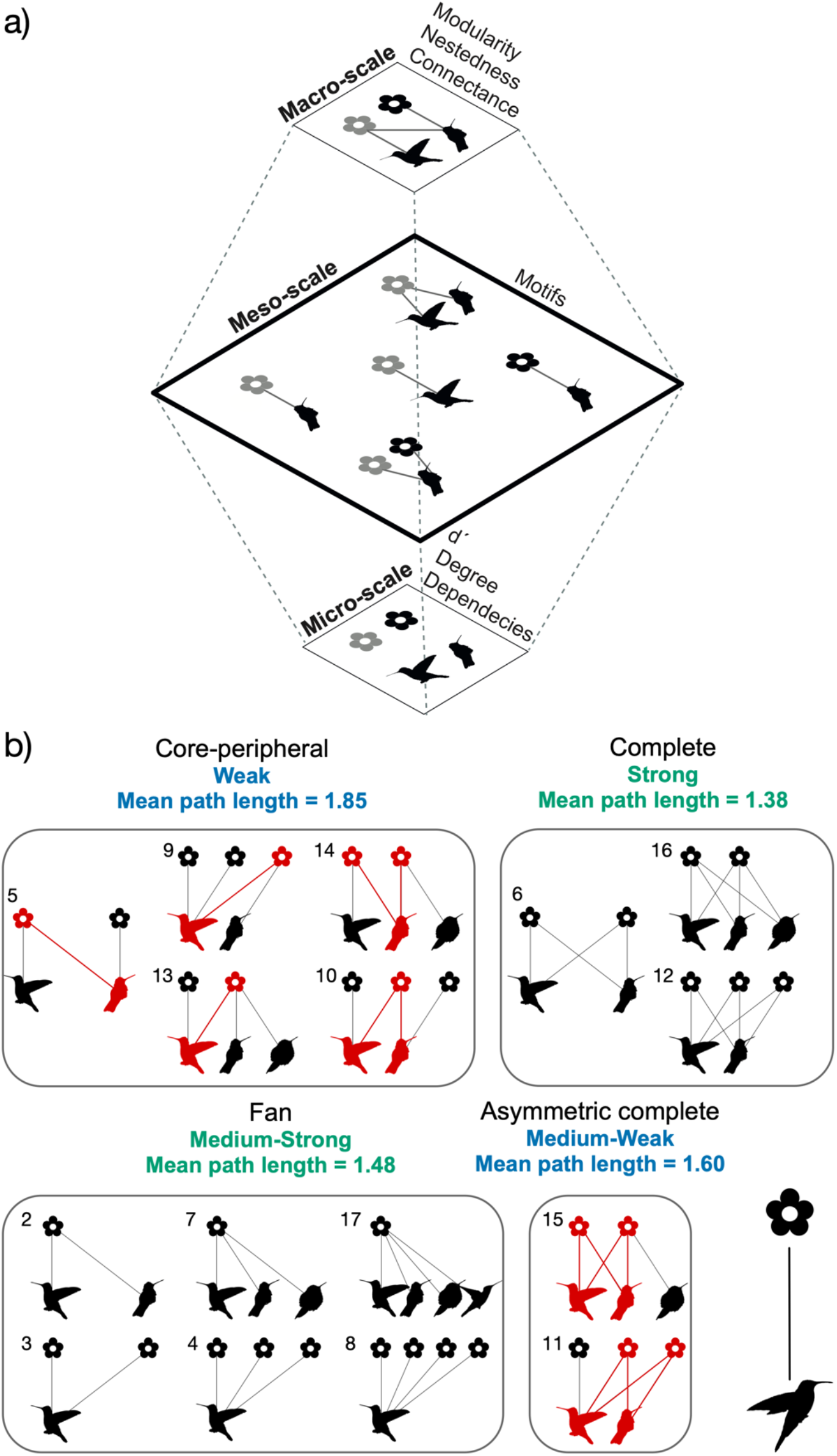
(a) Schematic showing how a small four-species network (top) can be characterised at three scales. Macro-scale metrics, such as modularity, nestedness and connectance, summarise the structure of the whole network. Micro-scale metrics, such as *d*’, degree or dependencies, characterise the structure of a single node. Motifs sit between these two extremes, at the meso-scale, capturing local-scale patterns of indirect interactions between species. The ‘meso-scale’ level shows the five types of motif that make up the macro-scale network. Note that the network itself is a four-species motif and so, for this example, we only consider motifs with fewer than four species (two- and three-species motifs). Importantly, motifs do not discard information about macro-scale structure. (b) A possible grouping of three- to five-node motifs by the broad indirect interaction structures they represent. Nodes in the bottom level of motifs are hummingbirds and nodes in the top level of motifs are plants. Each node represents a different species of animal or plant. Small numbers next to each motif are the ID of that motif, following Simmons, Cirtwill, et al. (2019) ‘Core-peripheral’ motifs contain a core of interacting generalist plants and pollinators (highlighted in red), connected to two or more peripheral specialist species. ‘Complete’ motifs are where generalists interact with generalists and all possible interactions are realised. ‘Fan’ motifs feature two or more specialists interacting with a single generalist. ‘Asymmetric complete’ motifs are the same as ‘complete’ motifs but linked to a single specialist. Thus, they are a particular type of ‘core-peripheral’ motif. Again, the core of interacting generalists is highlighted in red. For each motif, we calculated the mean path length (mean number of links between all pairs of nodes), and report the mean of these values across all motifs in each group below the group name. ‘Weak’, ‘Strong’, ‘Medium-Strong’, and ‘Medium-Weak’ indicate the possible strength of indirect interactions within motifs of each group. The strength of the indirect effect between two nodes tends to decay with increase path length between the nodes, such that nodes that are close to each other, topologically, likely have stronger indirect effects between them than nodes that are far apart. Thus, for example, nodes in ‘core-peripheral’ motifs with a high average path length likely experience weaker indirect effects than ‘complete’ motifs, which have short average path lengths.

As there is only a finite number of ways to arrange interactions between a given number of species, there is also only a finite number of motifs with a given number of nodes. In other words, all networks are made up of a limited number of different types of building block. For example, there are only 17 possible ways to arrange interactions between up to five species, and hence there exist 17 different motifs containing between two and five nodes (Figure 1b; note here that only 16 motifs are shown because we omit the simple two-node motif comprising a single link between one plant and one pollinator because it represents a direct interaction only without indirect effects).

In Figure 1b we propose a classification of motifs into four groups based on their indirect interaction structure. ‘Core-peripheral’ motifs are motifs comprising a core of interconnected generalist species, attached to two or more peripheral specialists. The mean path length between species in these motifs is high. *Path length* is defined as the number of links between two nodes. For example, in ‘core-peripheral’ motif 5, the path length between the black hummingbird and the black plant is 3 because the shortest (and in this case, only) path between these species involves 3 links (Figure 1b). The *mean path length* of a network is the average of the shortest paths between all pairs of species. The relatively high mean path length of 1.85 in core-peripheral motifs means that, on average, nodes are further apart. In turn, we expect indirect interactions in these motifs to be weaker, because indirect effects are expected to decay with increasing path length. For example, a change in the abundance of species *A* is likely to more strongly affect the abundance of species *B* if there is only one intermediary species (*A* → *C* → *B*) than if there are four intermediary species (*A* → *C* → *D* → *E* → *F* → *B*). As well as pathways between non-interacting species being longer, core-peripheral motifs also have fewer pathways between non-interacting species, because these motifs contain few links. Again, this reduces indirect effects (Guimarães et al., 2017) which, in turn, could help stop perturbations spreading through a network (Thébault & Fontaine, 2010).

‘Complete’ motifs stand in stark contrast to core-peripheral motifs. In complete motifs, all species interact with each other, creating many pathways between non-interacting species, and thus many ways for indirect effects to be transmitted. The mean path length is short, so the many indirect effects are also likely to be stronger. Predicting dynamics in complete motifs is likely to be harder, due to the multitude of possible pathways.

‘Asymmetric complete’ motifs are similar to complete motifs, except that one of the species in the motif is a specialist. Thus, asymmetric complete motifs contain a core where all species interact with each other, attached to a lone specialist species. Asymmetric complete motifs are a special case of ‘core-peripheral’ motif, with lower mean path length and higher number of pathways than the main set of core-peripheral motifs, and thus slightly stronger indirect effects. As the generalists in these motifs might be able to buffer changes in each other’s abundances, it is likely that the generalists have a stronger effect on the specialist, than *vice versa* (Simmons, Cirtwill, et al., 2019). The specialist species’ generalist partner has high levels of redundancy in its interactions and thus may be a reliable partner for the specialist. However, asymmetric complete motifs are likely less effective than core-peripheral motifs at curbing the spread of perturbations through the network as a whole, as most of their constituent species are involved in the hyper-connected core (Vieira & Almeida-Neto, 2015).

‘Fan’ motifs are the final group, comprising two or more specialists indirectly interacting via a shared generalist. These motifs extend the classic apparent competition and exploitative competition motifs from food webs to having any number of specialists interacting with a single generalist. Consider a motif where two plant species interact indirectly through a single pollinator species (the ‘fan’ motif 3 in Figure 1b). This could represent indirect facilitation, where an increase in the abundance of the first plant, increases the abundance of the shared pollinator which, in turn, increases the abundance of the second plant (Moeller, 2004; Sotomayor & Lortie, 2015). Alternatively, such a motif could represent exploitative competition for the pollinator or interference competition through heterospecific pollen deposition (Chittka & Schürkens, 2001; Flanagan, Mitchell, & Karron, 2010; Hochkirch, Mertes, & Rautenberg, 2012; Mitchell et al., 2009; Moeller, 2004; Simmons, Cirtwill, et al., 2019; Ye et al., 2014). Indirect interactions in fan motifs are relatively strong, with a medium-short path length.

By breaking down a network into its constituent motifs, it is possible to explicitly characterise indirect interaction structures between small groups of species, without losing any information about broader network structure. Specifically, motifs capture the topology of interaction chains, where changes in the abundance of one species influence the abundance of another species, through altering the abundance of one or more intermediary species (Simmons, Cirtwill, et al., 2019; Wootton, 1994). Even a simple four-species motif contains six different indirect interaction chains with up to two intermediary species (Figure 2). Larger and more complex motifs contain even richer detail on indirect interaction structures. This high level of detail is the advantage of the motif approach, allowing information about indirect interactions to be captured with a level of precision that is not possible when using traditional network metrics. Importantly, this extra information translates into novel and important insights into empirical data. For example, a recent study quantified species roles using a popular specialisation metric, *d’*, which measures the extent to which species’ interactions diverge from what would be expected if available partners were visited randomly (Kaiser-Bunbury et al., 2017). Using this metric, two key pollinator species were found to play similar roles in the community, both being super-generalists (Kaiser-Bunbury et al., 2017). However, when their roles were quantified using motifs, details of their indirect interactions were uncovered, revealing that the species actually played significantly different roles in the community: one was found to interact indirectly with generalist pollinators, while the other interacted indirectly with more specialist pollinators via shared specialist plants (Simmons, Cirtwill, et al., 2019).

**Figure 2:**
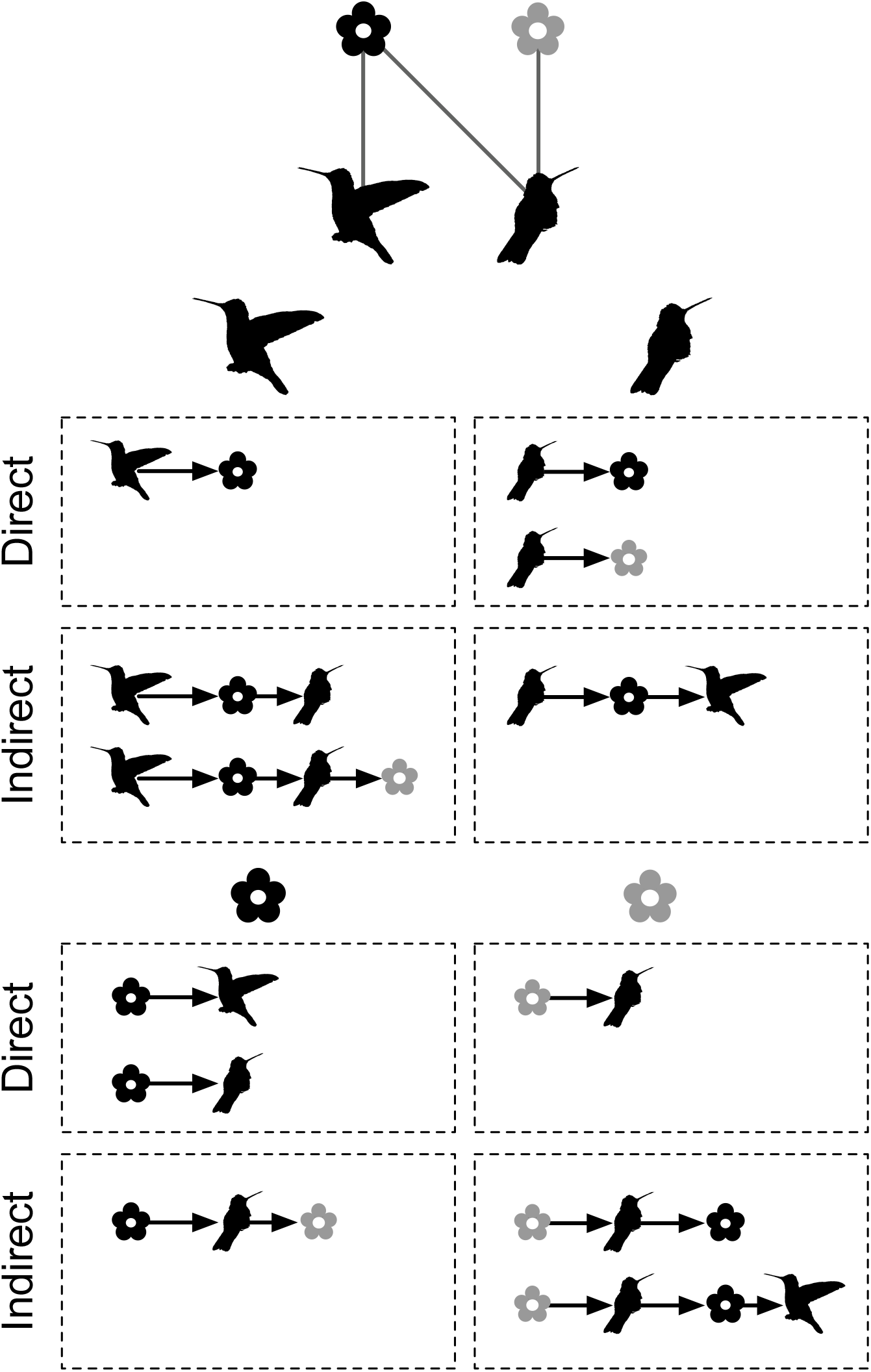
Motifs capture indirect interaction chains, defined as chains of species where a change in the abundance of one species alters the abundance of another species, through altering the abundance of one or more intermediary species (Simmons, Cirtwill, et al., 2019; Wootton, 1994). Here we consider a simple four-species motif and its constituent direct interactions and indirect interaction chains. It is clear how even a small motif contains rich detail on indirect interactions. The arrow shows the direction of the effect. For example, the left hummingbird has a direct interaction with the black plant, but also indirect interactions with the right hummingbird (via changing the abundance of the black plant) and the grey plant (via changing the abundance of the black plant and the right hummingbird).

We therefore used motifs to characterise indirect interactions in our analysis. Specifically, we calculated the mean frequency of all motifs up to five nodes (see motif topologies in Figure 1b) for each network and assembly process using the ‘bmotif’ R package (Simmons, Sweering, et al., 2019). To control for variation in network size, motif frequencies were normalised as a proportion of the total number of motifs within each motif size class (the number of nodes a motif contains) (Baker, Kaartinen, Roslin, & Stouffer, 2015). This was done to control for the fact that smaller motifs can be nested within larger motifs. As there is only one two-node motif (a single link between two nodes), and thus only one motif in the two-node size class, this was excluded from analyses because its normalised frequency would always equal one. Six-node, and larger, motifs were excluded because commonly-studied indirect interactions, like apparent competition, are present in smaller motifs and five-node motifs already contain varied and long interaction chains with up to three intermediary species between two indirectly interacting partners. Limiting to five-node motifs was also beneficial for visualisation, interpretation and computational reasons.

### Statistical analysis

We used an ANOVA framework to assess statistical differences between the frequencies of motifs in networks generated using different assembly processes. First, a MANOVA was used with frequencies of all 16 motifs as dependent variables and assembly process as the independent variable to determine whether there was an overall effect of assembly process on motif frequency distribution. Then, to identify how assembly processes affect specific dependent variables, we conducted univariate ANOVAs for each motif. For this, pairwise comparisons between assembly processes were calculated using the ‘multcomp’ R package (Hothorn, Bretz, & Westfall, 2008). Adjusted p-values were used to account for multiple comparisons, using the ‘single-step’ method in ‘multcomp’.

## Results

Different assembly processes produced significantly different motif distributions (MANOVA: df = 4, F = 2530.5, *p* < 0.001): neutral processes (abundance) were associated with more occurrences of complete, asymmetric complete and fan motifs (motifs 6, 8, 11, 12, 16 and 17), while niche-based processes (morphological match and phenological overlap) were associated with more occurrences of core-peripheral motifs (motifs 5, 10 and 14) (Figures 1b, 3 and 4). Furthermore, some differences were observed between morphological matching and phenological overlap matrices: phenological overlap matrices had significantly higher frequencies of motif 9 (a core-peripheral motif) than morphological matching, but significantly lower frequencies of motif 14 (another core-peripheral motif; Figures 1b, 3 and 4).

**Figure 3:**
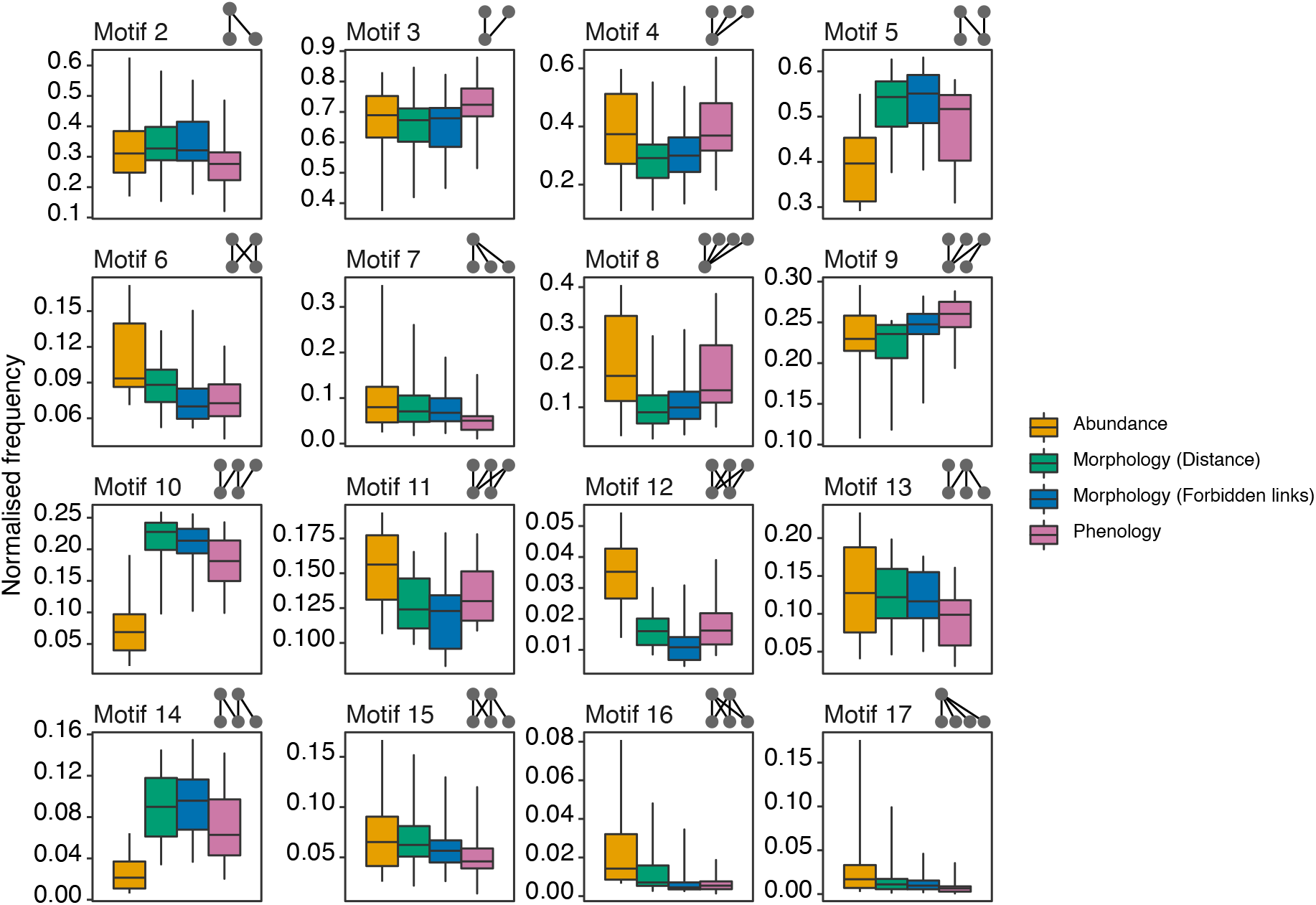
Normalised frequencies of each motif for networks generated using abundance (neutral processes) and morphological matching and phenological overlap (niche-based processes) for 24 plant-hummingbird interaction networks sampled across the Americas. Boxplots represent the distribution of mean normalised motif frequencies for generated networks across the 24 sets of abundance, morphology and phenological data. Upper whiskers represent 95% quantiles, the upper hinge is the 75% quantile, the middle line is the median, the lower hinge is the 25% quantile and the lower whisker is the 5% quantile. See Figure 4 for significance levels. In the motifs depicted above each boxplot, nodes in the bottom level of motifs are hummingbirds and nodes in the top level of motifs are plants.

**Figure 4:**
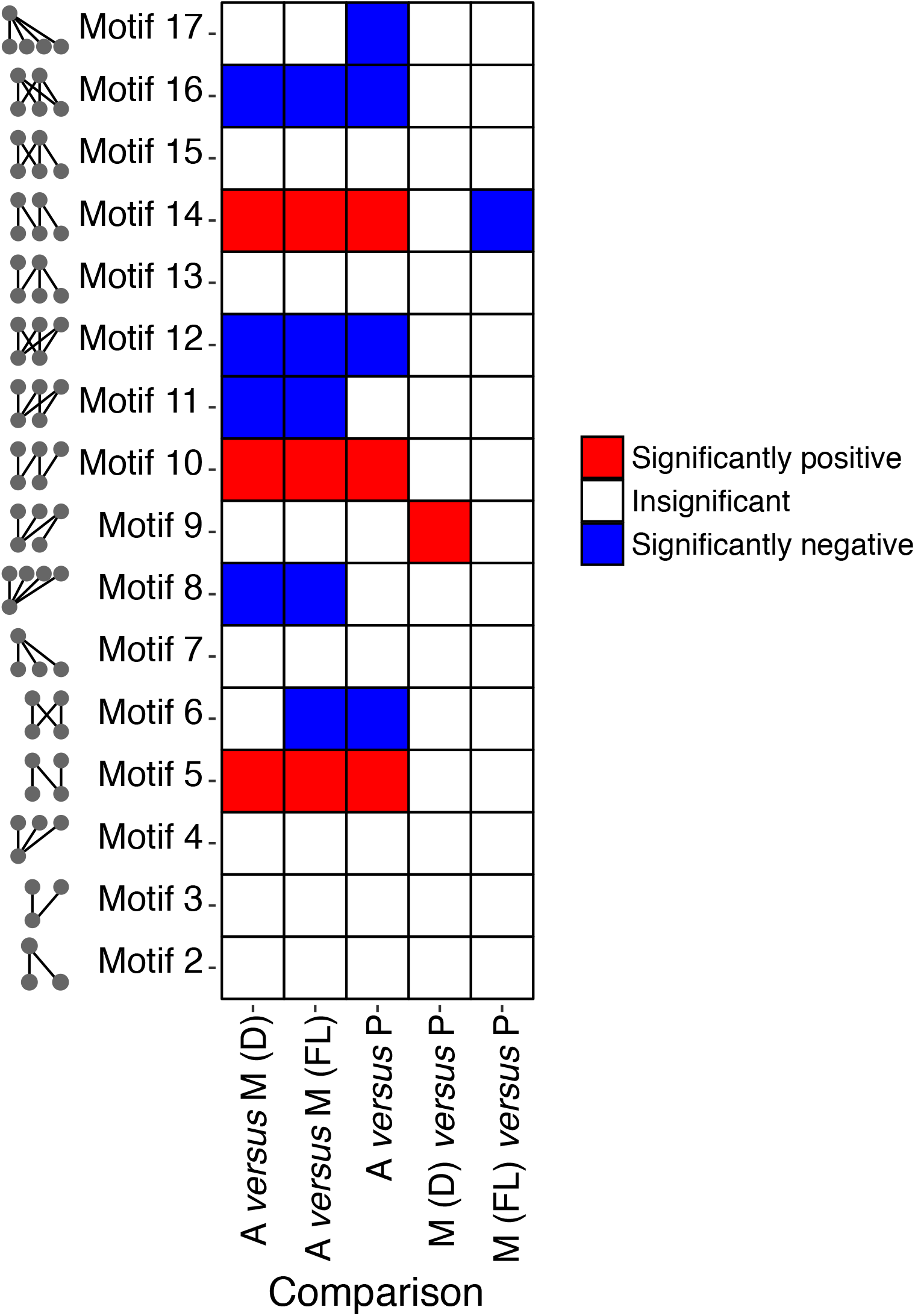
Matrix showing whether there are significant (adjusted *P* < 0.05) differences in normalised motif frequencies depending on the assembly processes (neutral processes like abundance, or niche-based processes like morphological matching or phenological overlap). Abbreviations for assembly processes are: ‘A’ is abundance, ‘M (D)’ is morphological matching based on distance between corolla depth and bill length, ‘M (FL)’ is morphological matching based on the forbidden link concept, and ‘P’ is phenology. Comparisons are relative to the first processes expressed. For example, if a cell in the A *versus* M (D) column is red, this means the motif frequency was significantly higher in the M (D) matrices than in the A matrices. Conversely, if a cell in the A versus M (D) column is blue, this means the motif frequency was significantly lower in the M (D) matrices than in the A matrices.

## Discussion

We find that networks generated using different assembly processes have significantly different patterns of indirect interactions. Networks governed by neutral effects (species abundance) tend to have more densely-connected complete, asymmetric complete and fan motifs where either (i) indirect interactions between plants/pollinators are mediated through a single pollinator/plant (fan motifs 8 and 17), or (ii) indirect interactions may be strong because there are multiple routes for indirect effects to travel at the same time (complete and asymmetric complete motifs 6, 11, 12 and 16) (Figures 3 and 4). Conversely, networks produced assuming niche-based processes – those determined by morphology or phenology – contain more core-peripheral motifs, that comprise a core of interacting generalists, supported by peripheral specialists (core-peripheral motifs 5, 10, 14) (Figures 3 and 4).

Neutral processes produced two main types of motifs. First, they produced motifs, where specialists affect each other indirectly via a single generalist (fan motifs, such as motifs 8 and 17). These fan motifs (Simmons, Cirtwill, et al., 2019) extend the classic apparent competition and exploitative competition structures from food webs (motifs 2 and 3) to having more than two specialists. Importantly, despite being generated by the same process, motifs 8 and 17 are likely to have different levels of competition between the specialist species. In motif 8, many plants compete for a single pollinator (Figure 1b). In this situation, competition is likely to be low between the plants, especially if the pollinator is abundant, as the plants only need one successful visit from a pollinator to disperse their pollen and reproduce. Conversely, in motif 17, multiple pollinators are competing for a single plant (Figure 1b). Here, competition is likely to be stronger, as pollinators are relying on the plant as a regular, limited food source. Importantly, however, these networks represent mutualistic interactions between species and thus it is also possible that ‘fan motifs’ represent indirect facilitative, rather than competitive, situations, where specialists indirectly benefit each other through interactions with a single generalist (Moeller, 2004; Sotomayor & Lortie, 2015). For example, the presence of a plant species could increase pollinator visits to one or more coflowering species, or multiple plant species could combine to form a large, shared floral display that increases pollinator visitation to all coflowering plants beyond what would be expected if each of the plants flowered in isolation. Whether indirect interactions are competitive or facilitative can depend on a range of factors, such as the distance between plants and their spatial configuration (Charlebois & Sargent, 2017), however there is evidence that pollinator abundance can have an influence, with facilitation occurring above a threshold abundance, and competition occurring below the threshold (Ye et al., 2014). Thus, combining motif analysis with information on empirical or simulated population dynamics, could give insight into the directionality of indirect effects.

The second type of motif produced by neutral processes is complete and asymmetric complete motifs which have many links, providing many possibly pathways through which indirect effects can flow (motifs 6, 11, 12, 16). This likely results from the neutral model’s lack of consideration of ‘forbidden links’ (Canard et al., 2014; Jordano, 2016b): as long as two species are of sufficiently high abundance, they are able to interact, resulting in more pathways (Simmons et al., 2018). This is in contrast to niche-based processes, where poor morphological matches or low temporal co-occurrence would prevent some interactions from being formed. This has important implications for whole-network dynamics, as it suggests that under neutral processes, the average length of the interaction chain between any two species will be lower, increasing the magnitude and number of indirect effects, but decreasing their localisation. In turn this could allow the spread of perturbations through the community (Thébault & Fontaine, 2010). In complete motifs 6, 12 and 16, all plants interact with all pollinators. Here we might expect indirect interactions to be strong, as effects can be transmitted through multiple links simultaneously and the indirect interaction chains are shorter, but also less predictable (Simmons, Cirtwill, et al., 2019). For example, in motif 12, if a pollinator decreased in abundance, this would remove the mutualistic benefit to the three plant species, but could also reduce competition between the two pollinators (Simmons, Cirtwill, et al., 2019); further research is necessary to examine the complex dynamics that could occur in these motifs. Motif 11 represents a slightly different situation to that in 6, 12 and 16, as motif 11 has a single specialist interacting with a completely connected set of generalists. This is therefore an asymmetric complete motif, where it has been suggested that generalists have a stronger effect on the specialists than the specialists have on the generalists, as the generalists are able to buffer changes in each other’s abundances (Simmons, Cirtwill, et al., 2019).

Niche-based processes resulted in motifs with a core of interacting generalists, connected to peripheral specialists (core-peripheral motifs 5, 10, 14). The indirect interaction pathways in these motifs can be highly complex. For example, in motif 5, there are four species: two plants in the top left (P_L_) and top right (P_R_), and two hummingbirds in the bottom left (H_L_) and bottom right (H_R_). One possible pathway is that P_L_ can negatively affect H_R_ indirectly, by providing a mutualistic benefit to H_R_’s competitor H_L_, and by competing with P_R_, reducing the mutualistic benefit to H_R_ (Simmons, Cirtwill, et al., 2019; Vázquez, Ramos-Jiliberto, Urbani, & Valdovinos, 2015). While a complete study of the dynamics of each motif is beyond the scope of this work, our results do suggest that niche-based processes restrict the sharing of interaction partners, thus forcing indirect pathways between species to be longer. Given that longer pathways likely have weaker indirect effects, niche-based processes likely reduce the magnitude of indirect effects in the community (Guimarães et al., 2017). In turn, this could limit the spread of perturbations through the network (Thébault & Fontaine, 2010).

While there were few differences between different niche-based processes, networks based on phenological overlap had significantly higher frequencies of motif 9 (a core-peripheral motif with two generalists interacting) and significantly lower frequencies of motif 14 (a core-peripheral motif with three generalists interacting) than morphological matching models. This could reflect the degree of constraint between these two processes. *A priori*, it is difficult to say whether phenological overlap or morphological matching represents a greater constraint on species interactions. Phenological overlap requires species to co-occur in time to interact, but ignores species morphology, while morphological matching only allows species to interact if the hummingbird bill length and floral corolla depth are sufficiently matching, regardless of temporal co-occurrence. For our data, the phenological overlap model produced significantly more motifs with two generalists, and significantly fewer motifs with three generalists, than the morphological matching model. This suggests that interactions between generalists are rarer under phenological overlap, indicating that lack of phenological overlap may impose more forbidden links than morphological mismatch in plant-hummingbird pollination systems.

Here we shed light on the different processes associated with patterns of indirect interactions in mutualistic networks, quantified using motifs. As well as being ecological interesting in its own right, our results are also useful for interpreting the results of motif analyses and for generating motif-driven hypotheses. For example, networks with a high proportion of invasive species may be expected to have higher frequencies of motifs associated with neutral effects, because invasive species lack the coevolutionary associations of native species (Jeferson Vizentin-Bugoni et al., 2019). Overall, our results link indirect interaction structures to distinct generative processes. The normalised motif profiles we present represent a baseline of what structures would be expected in communities dominated by morphological matching, phenological overlap or neutral effects. By measuring the similarity of motif profiles from empirical networks to those idealised profiles presented here, it may be possible to infer the processes acting in a community from the indirect interaction topology alone, and thus inform the type of conservation actions that are needed. Further research along this line is necessary, alongside empirical validation, but our findings suggest potential for using structure as a proxy for processes in a conservation context.

## Acknowledgements

BIS and APB are supported by the Natural Environment Research Council [NE/S001395/1]. BIS is also supported by a Royal Commission for the Exhibition of 1851 Research Fellowship. BD thanks the Danish National Research Foundation for its support of the Centre for Macroecology, Evolution and Climate (grant no. DNRF96).

